# Healing helminths: The disease-modifying potential of helminth-derived proteins in animal models of inflammatory disease - a systematic review with cross-study quantitative analysis

**DOI:** 10.64898/2026.04.02.716049

**Authors:** Sienna Stucke, Aonghus Feeney, Richard Lalor, Sheila Donnelly, John Pius Dalton, Declan McKernan, Eilís Dowd

## Abstract

Helminths are parasitic worms that secrete a variety of immune-regulating molecules to modulate the host’s inflammatory responses, enabling them to persist within the host over a long period of time, even decades. Their capacity to control host responses has prompted research into helminth-derived molecules as potential therapies for controlling excessive immune and inflammatory activity across a range of diseases. This systematic review with cross-study quantitative analysis aims to synthesize the published data on helminth-derived immunomodulatory peptides/polypeptides/proteins (HDIPs) with a focus on determining the extent of their disease-modifying and anti-inflammatory potential in *in vivo* animal models of inflammatory disease. In accordance with PRISMA 2020 guidelines, a predefined systematic search of the PubMed, Web of Science and Medline databases identified relevant studies published up to February 2026, and 65 articles were included after screening. We found that, although the HDIPs were assessed in multiple different disease models, most published studies assessed their potential in mouse models of colitis, asthma, arthritis and sepsis. Twenty species from which >65 isolated HDIPs were derived were tested in these models, with the trematode, *Fasciola hepatica*, and the nematode, *Acanthocheilonema viteae*, the most explored species. A common property of the HDIPs was the ability to significantly reduce disease severity across the *in vivo* animal models of inflammatory disease, underpinned by a decrease in pro-inflammatory cytokine levels and an increase in anti-inflammatory cytokine levels. Overall, this systematic review with cross-study quantitative analysis not only synthesizes the existing literature in this field but also highlights the disease-modifying and anti-inflammatory potential of HDIPs for a range of diseases in which immunoregulatory therapeutics may improve disease outcomes. It also encourages accelerated advancement of these helminth-derived molecules into first-in-human clinical trials.

## INTRODUCTION

Helminths are parasitic worms that infect both humans and animals and are among the most widespread infectious agents globally. This is mainly in developing regions with poor sanitation, where worms, particularly soil□transmitted helminths, are transmitted via contaminated water or food, or by contact with soil contaminated with human faeces containing helminth eggs. These infections contribute substantially to the global disease burden, particularly in tropical and subtropical areas where they cause chronic morbidity, nutritional impairment, anaemia, and developmental delays in children^1–3^. Soil□transmitted helminthiases alone affect an estimated 1.5□billion people worldwide, ranking among the most common neglected tropical diseases identified by the World Health Organization.

Helminths are able to persist within their human or animal hosts for months, years, or even decades by secreting a diverse array of molecules that help them evade the host’s immune system, thereby avoiding immune-mediated rejection^4^. These helminth-derived immunomodulatory peptides, polypeptides, and proteins (HDIPs) can influence the host immune system through multiple coordinated immunomodulatory mechanisms that suppress inflammation to promote their survival. This modulates the balance between Th1 and Th2 immune responses, leading to a reduction in pro-inflammatory Th1 cytokines and an increase in anti-inflammatory Th2 cytokines. Ultimately, worm-mediated immune modulation creates an immune environment that allows the helminth to establish and maintain long-term chronic infection^5^. Intriguingly, the immunomodulatory properties of helminths have positioned them as a double-edged sword. While helminths can appear as “foes” by contributing substantially to global disease burden, their capacity to dampen inflammation and modulate immune responses has prompted researchers to explore their therapeutic potential in diseases driven by overactive immunity^4,6^. Consequently, helminths are not merely pathogens to be eradicated; rather, as live organisms or through their HDIPs, they represent a promising “friend” by providing a novel biotherapeutic avenue for the treatment of autoimmune and inflammatory disorders.

To date, most research in helminth therapy has focussed on using live inoculation with helminth eggs or larvae across a variety of inflammatory and immune-mediated diseases. Clinical trials have primarily targeted conditions such as inflammatory bowel disease (Crohn’s disease and ulcerative colitis), celiac disease, allergic disorders including asthma and allergic rhinitis, and multiple sclerosis. But while early open-label studies demonstrated that live helminth inoculation is generally safe and well tolerated, subsequent randomised controlled trials have reported only modest or inconclusive clinical benefits, highlighting the need for larger, more rigorously designed studies to fully evaluate therapeutic efficacy^7^. Additional challenges are that the global pathogenic effect of live parasitic infections is unknown and potentially difficult to control, and that cultural attitudes and social stigma surrounding live parasite infections can foster public scepticism or uneasiness regarding helminth therapy^8^.

An alternative approach that has gained significant traction in recent years is the use of specific HDIPs as therapeutic molecules. Multiple different HDIPs from a variety of helminth species have been isolated, purified and characterised over the past twenty years. These have been consistently shown to module inflammatory responses in cellular and animal models of inflammatory disease, but to date, just one peptide - P28GST, a HDIP with glutathione S-transferase activity isolated from schistosomes – has progressed to early clinical trial^9^.

To support evidence-based evaluation of the immunomodulatory potential of HDIPs, we recently conducted a systematic review examining their anti-inflammatory effects in cellular models of inflammation^10^. However, no equivalent systematic synthesis has been performed for *in vivo* studies in animal models of inflammatory disease. Accordingly, this review aims to systematically collate and analyse published data on HDIPs in animal models, with a focus on assessing the extent of their disease-modifying and anti-inflammatory effects in physiological contexts.

## METHODS

### Search strategy

This study was completed in accordance with the PRISMA 2020 guidelines^11^ to find articles in which HDIPs were assessed in animal models of inflammatory disease. The literature search was completed in the PubMed, Web of Science and Medline databases with the specific search string: helminth AND secret* AND (immunomod* OR immunosup* OR antiinflam*). After removal of duplicates, this search identified a total of 829 records, spanning July 2001 to February 2026. These articles were then screened according to the strategy outlined below and depicted in the PRISMA flow diagram (Fig. 1). This yielded 65 articles which were included in this systematic review. All records were managed in the Endnote and Microsoft Excel software packages.

**Fig. 1.**
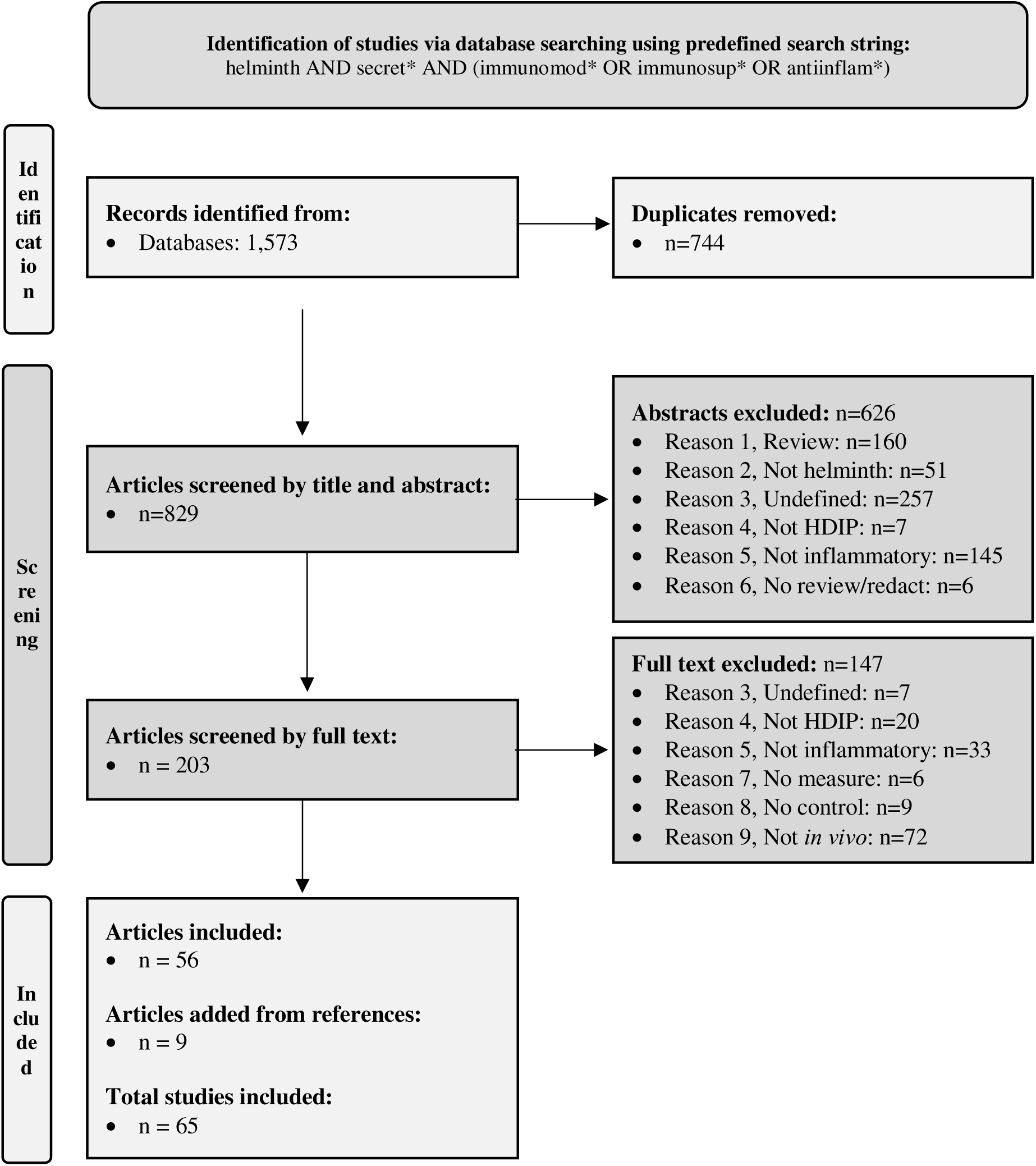
PRISMA flow chart. Flow chart based on PRISMA 2020 guidelines^11^ detailing the screening strategy employed for the study selection in the present study.

### Screening strategy

The remaining articles were screened by title and abstract according to the following inclusion and exclusion criteria. Inclusion criteria: (1) original research study, (2) HDIPs, (3) *in vivo* animal model of inflammatory disease. Exclusion criteria: (1) review article, (2) not helminth-derived (3), undefined helminth-derived molecule, (4) not HDIP, (5) not tested in a model of inflammatory disease and (6) not peer-reviewed or redacted. In the full-text screen, three other exclusion criteria were applied (7) no cytokine or disease measure, (8) no appropriate control and (9) not *in vivo*. After screening, 56 articles met the inclusion and exclusion criteria. The reference lists in these articles were then further screened by title, and those naming a specific worm species (rather than using the term “helminth”) were further screened using the same inclusion and exclusion criteria as the other articles. This led to inclusion of a further 9 articles. Thus, a total of 65 articles were included in this systematic review (Table 1 and Supplementary Excel file).

**Table 1.**
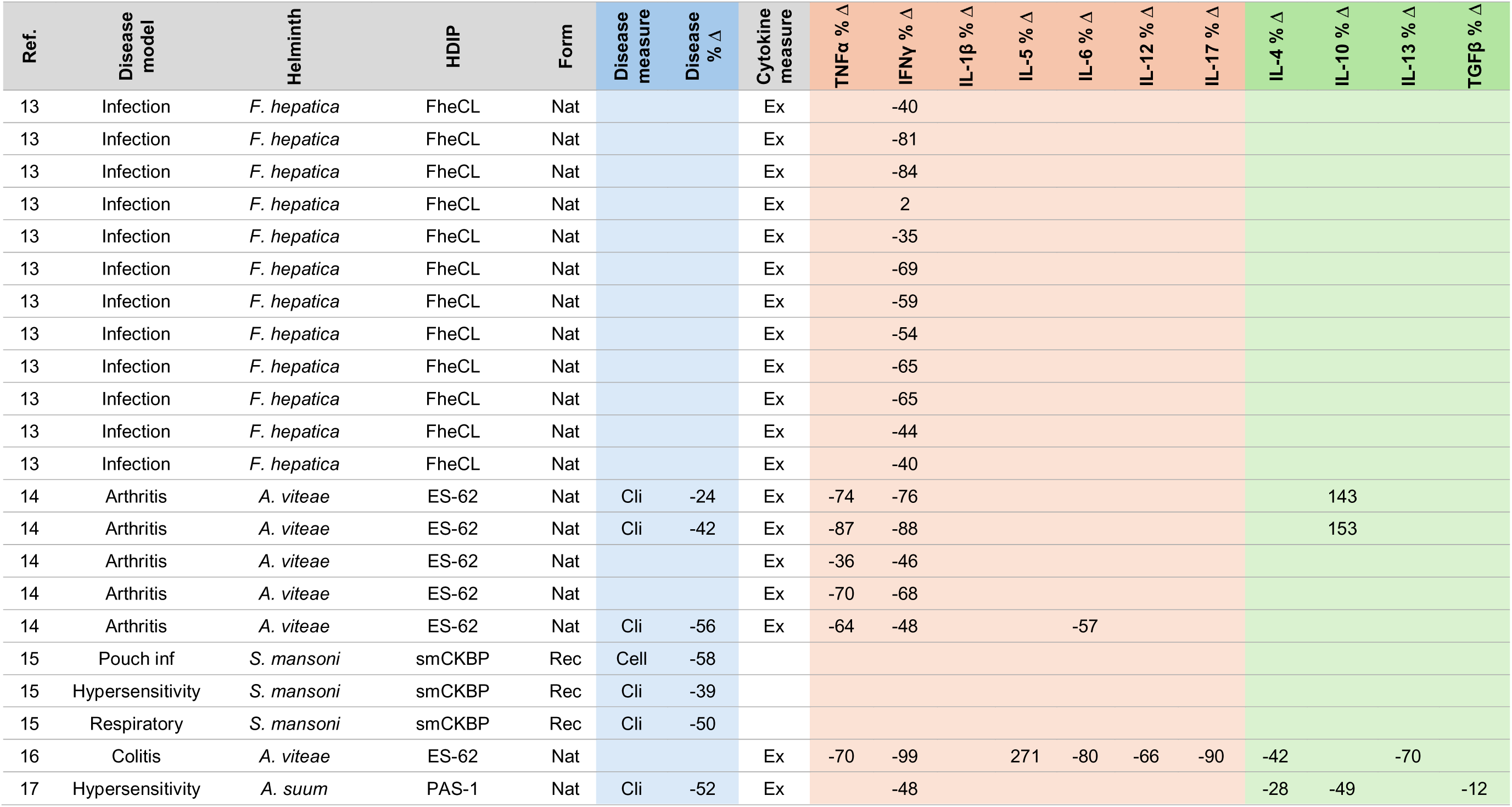
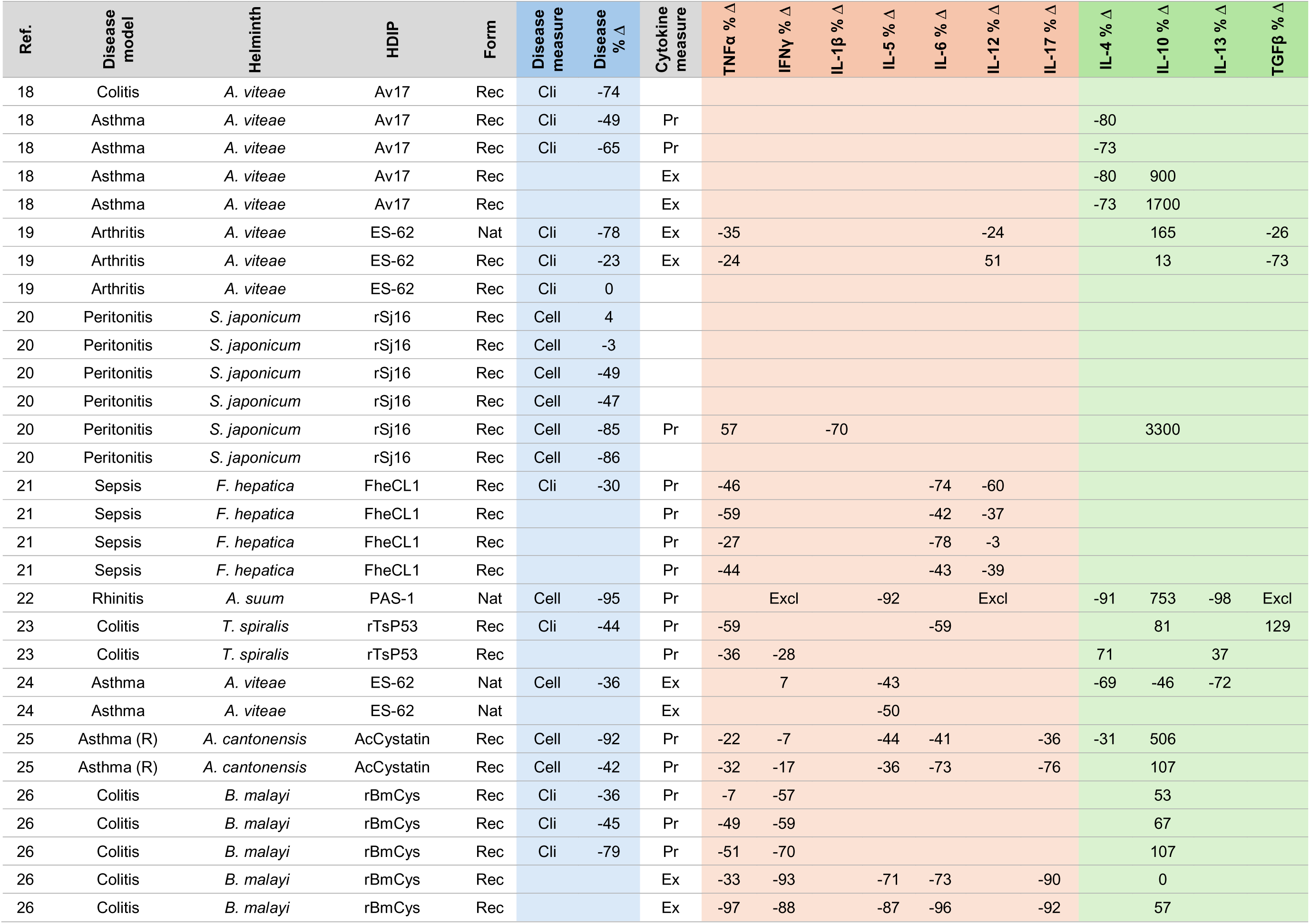
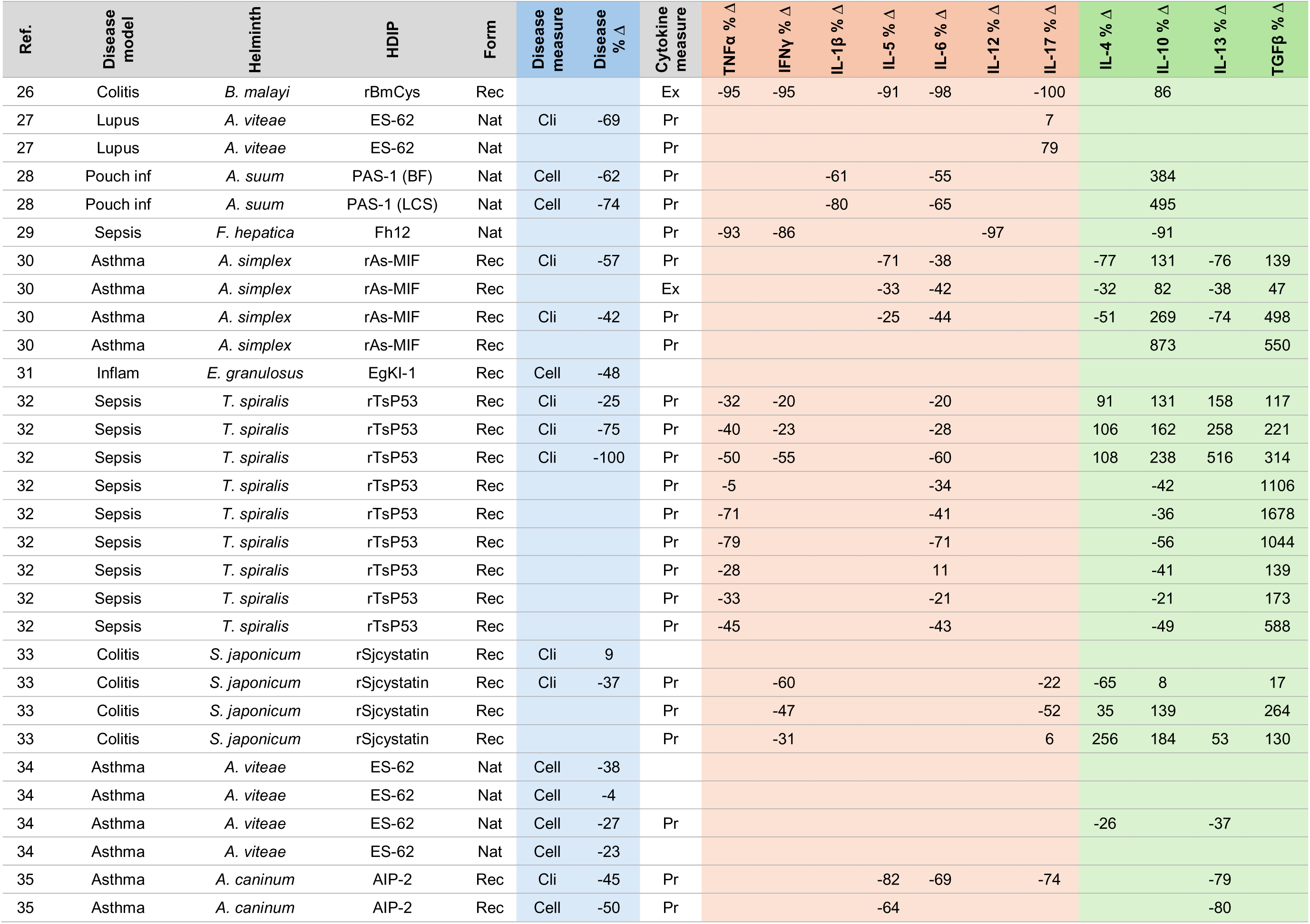
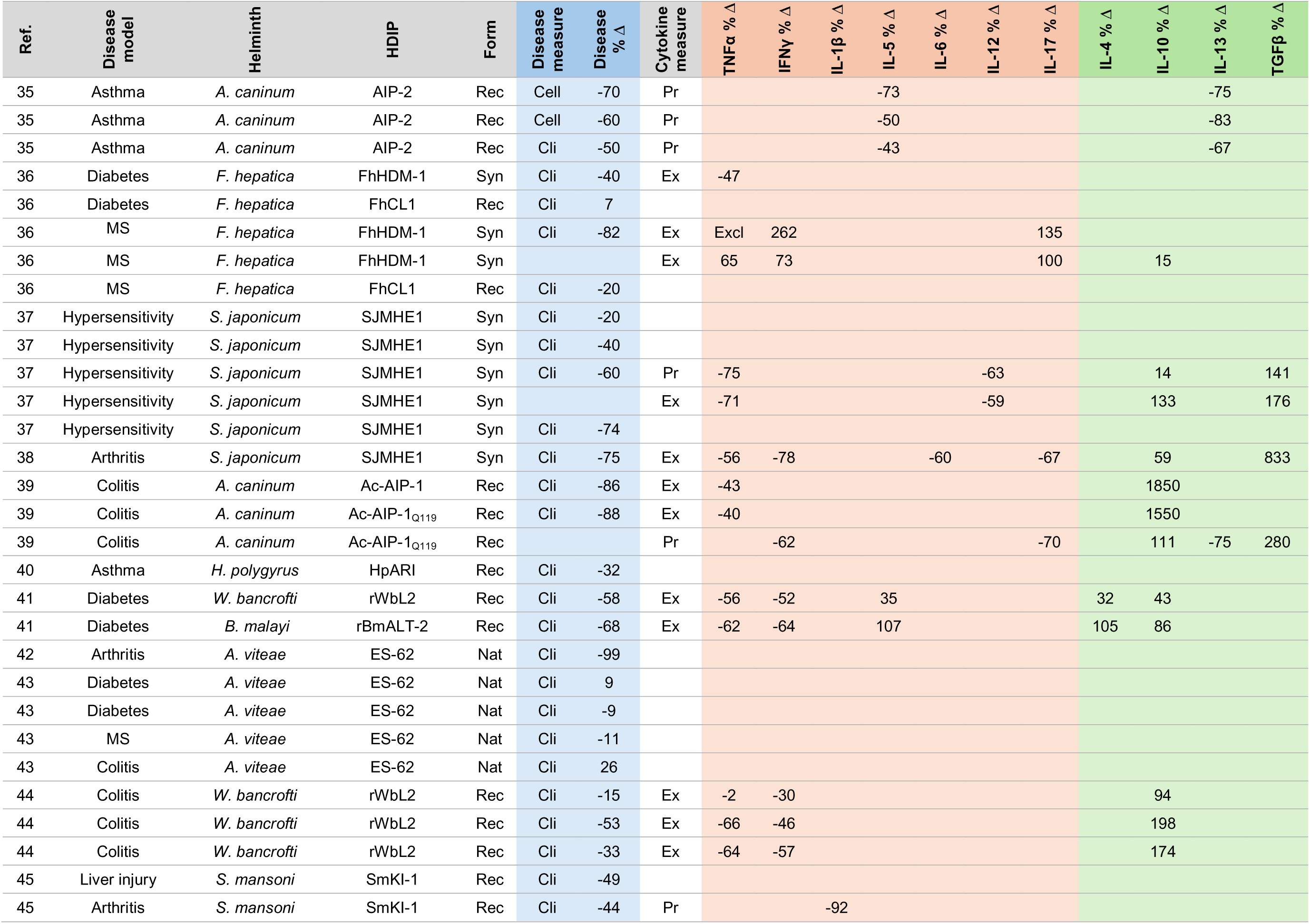
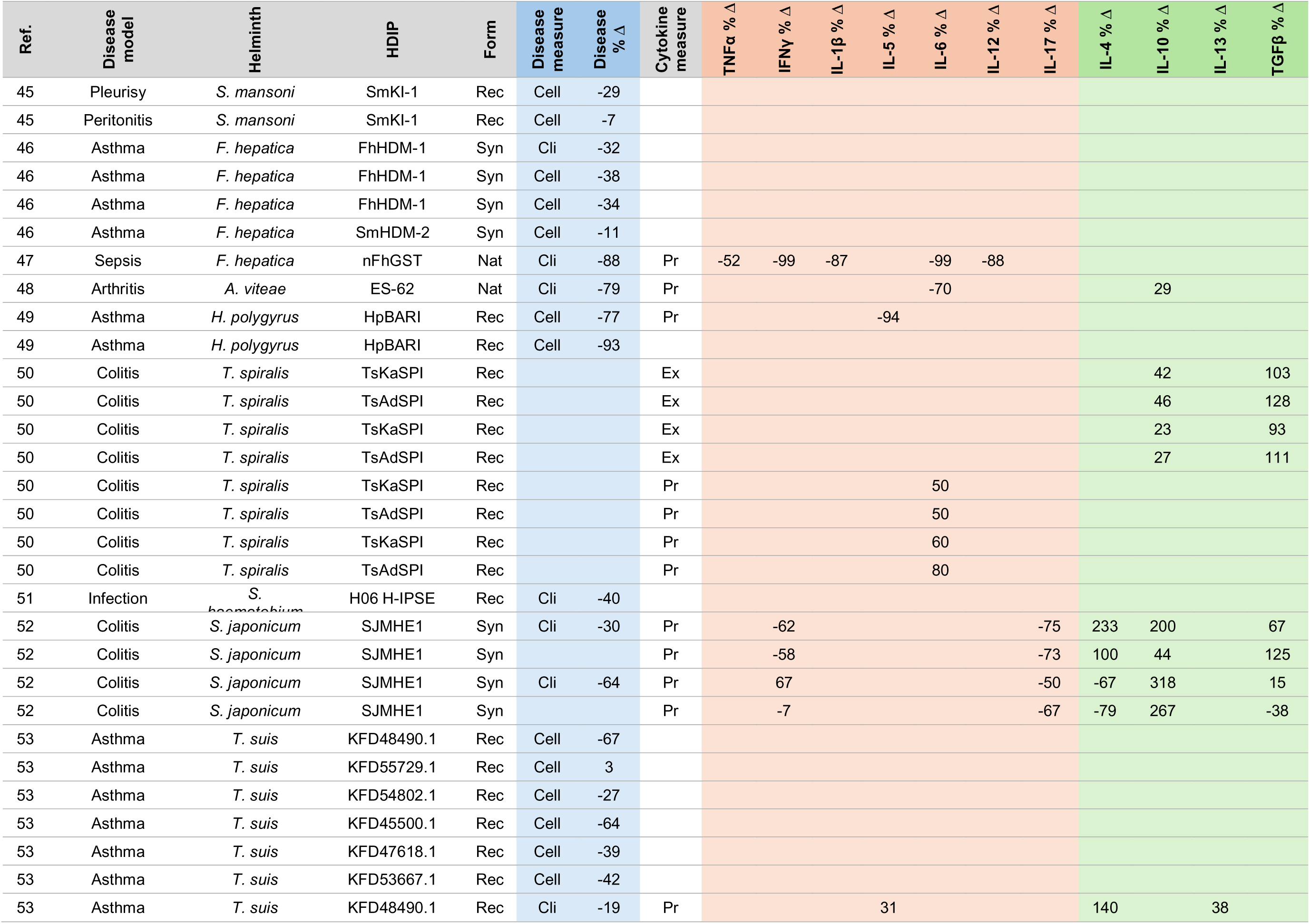
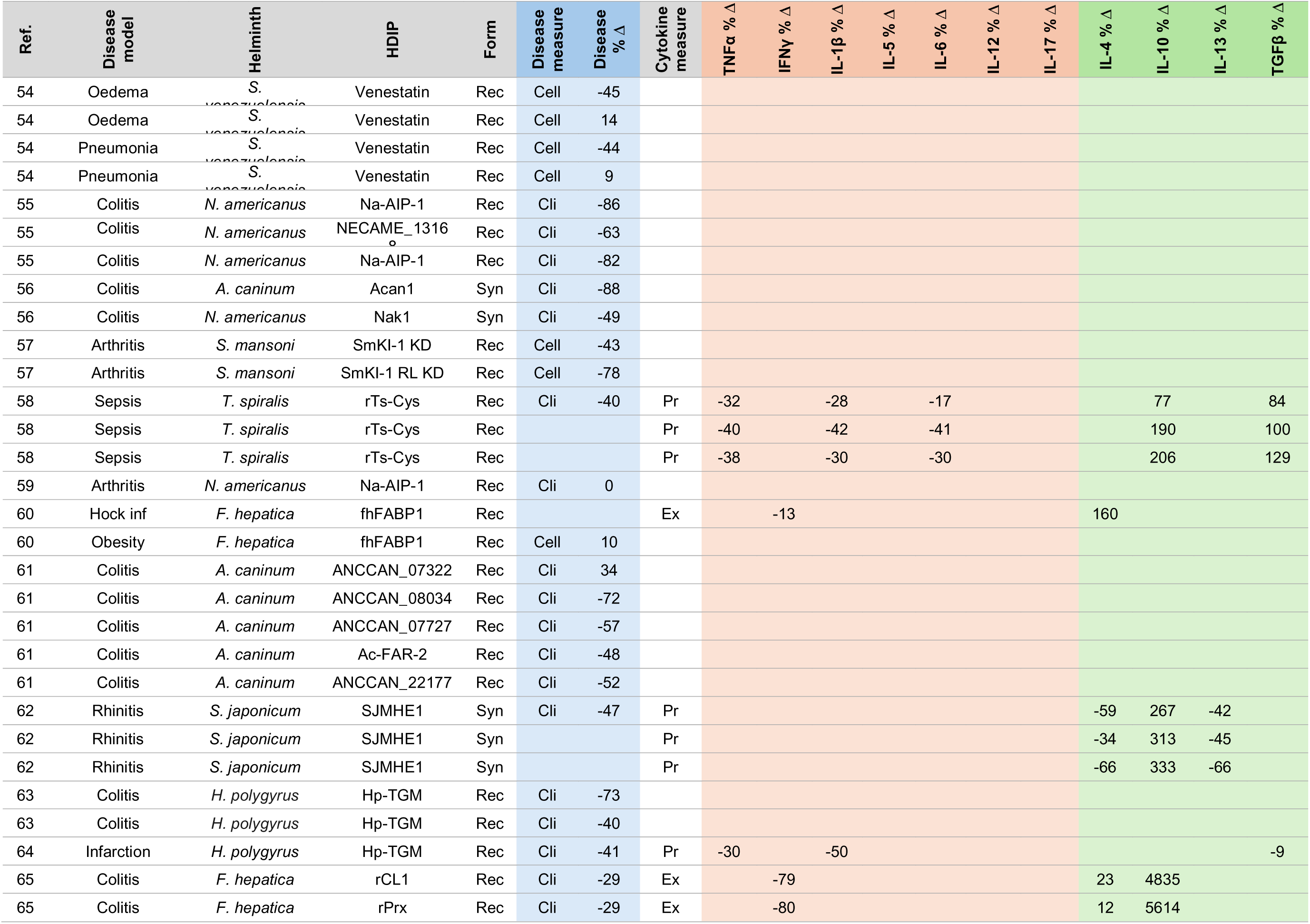
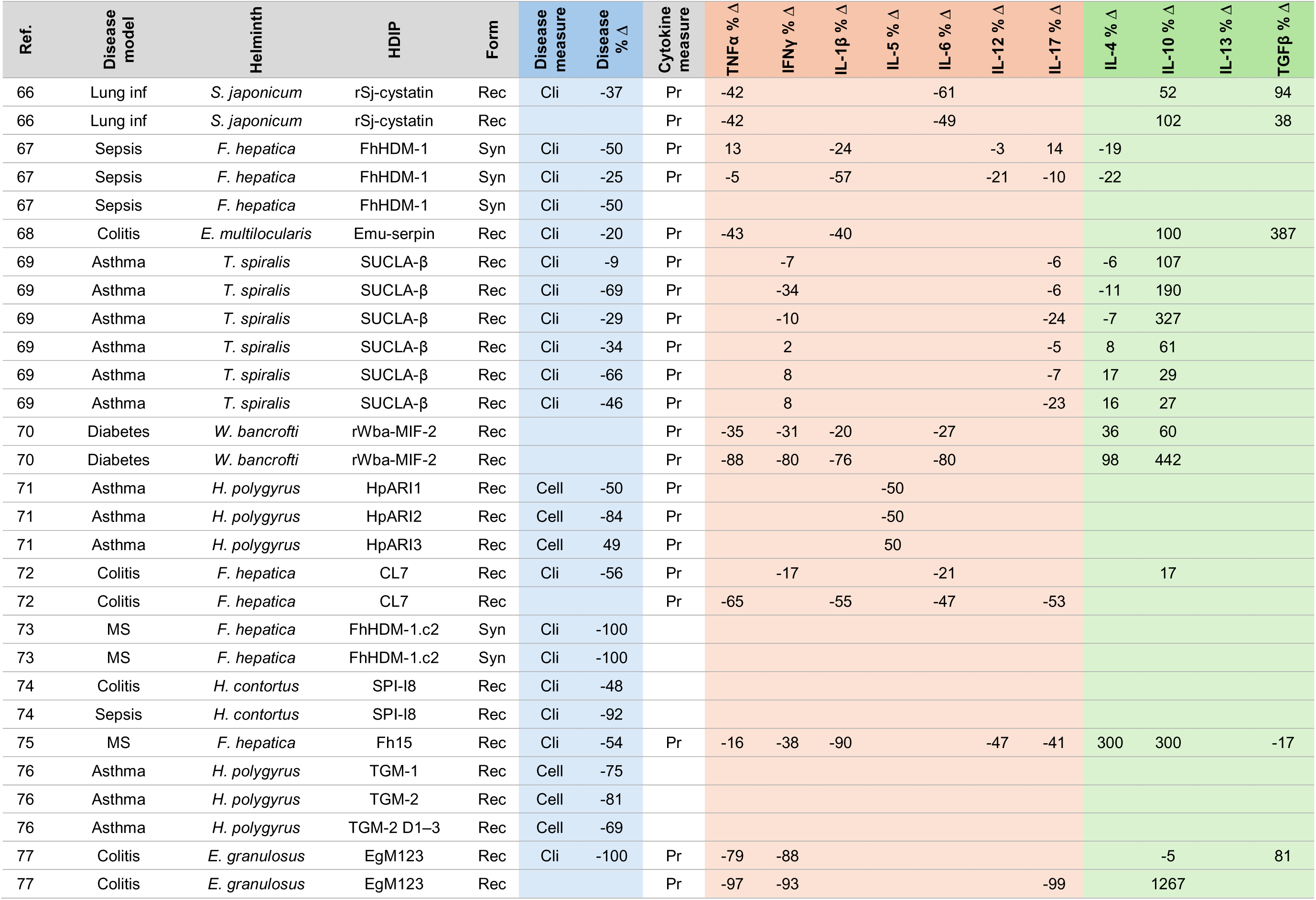
Studies and individual records included in this review. A total of 65 articles were included in this review from which 141 and 132 records were extracted in which the effects of the HDIPs were assessed on disease severity and cytokine levels respectively. For each of these records, the percent change in disease burden and cytokine levels in the animal models without and with HPID are shown. All diseases were modelled in mice with one exception denoted by (R). See Supplementary Excel file for more details and specific values. Abbreviations: MS: Multiple sclerosis; Inf: Inflammation; Nat: Native; Rec: Recombinant; Syn: Synthetic; Cli: Clinical; Cell: Cellular; Ex: *Ex vivo*; Pr: Primary; Excl: Values excluded as statistical outliers. Some cytokines were measured infrequently (<5 articles) and these were omitted from the review.

**Table 1.**
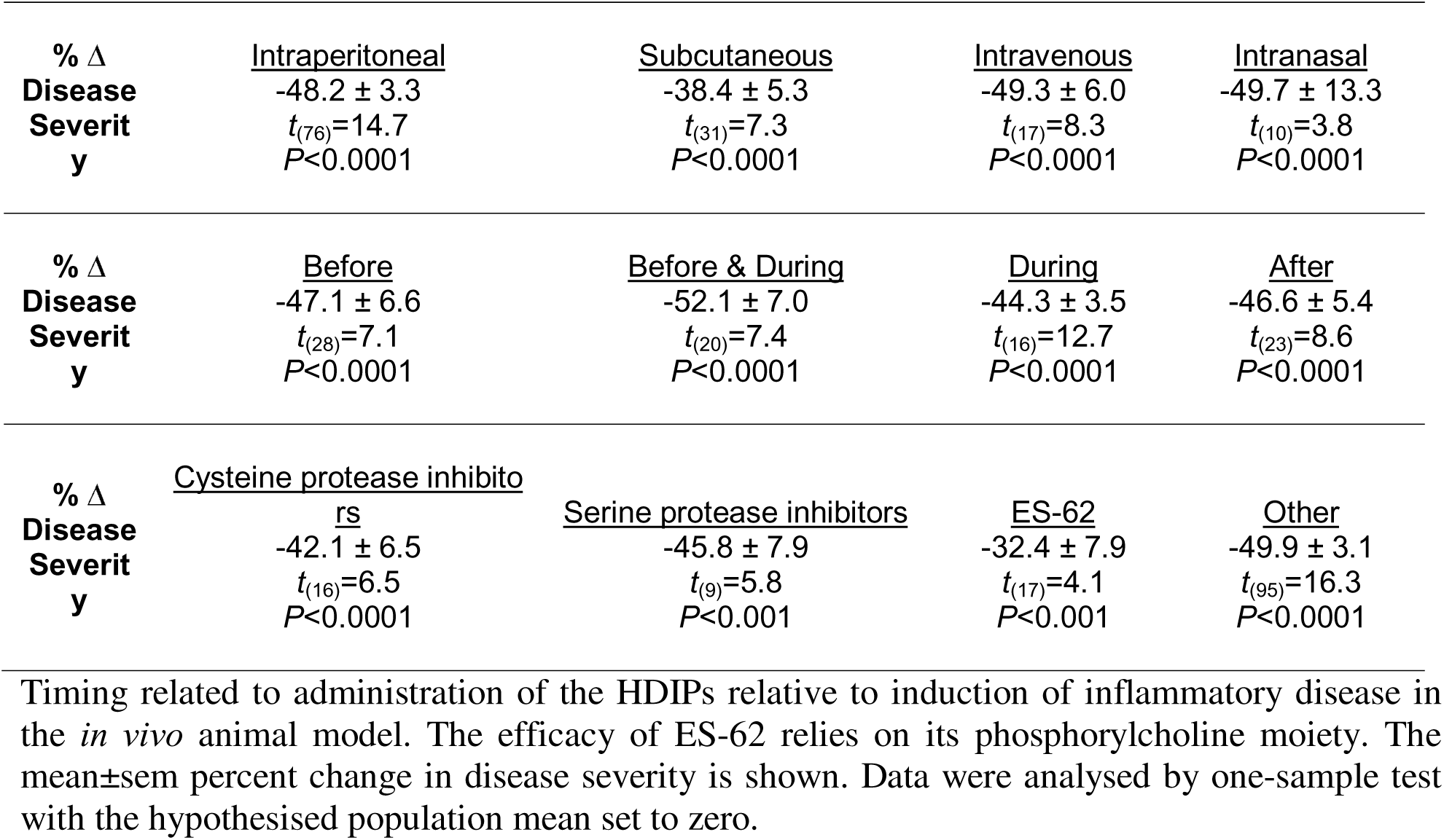
Effect of HDIPs on disease severity with different routes of administration, timing of administration and mechanistic classes.

### Variables extracted

Variables manually extracted from the selected articles included disease studied, disease model and species, helminth species, HDIP name and form, as well as disease burden and cytokines measured (Table 1 and Supplementary Excel file). Because any given article may have had several experimental parameters (e.g. different HDIP doses), to comprehensively synthesize this literature, each of these was considered a separate “record” for data analysis. For each record, a single measure of disease burden was extracted in studies where this was assessed. Either a clinical measure was taken (most commonly a disease score or index), or if this was not available, then a cellular measure was taken (most commonly a cell count). In the selected articles, cytokine levels were measured in primary sources taken from the animal models (tissues, cells, fluids) and/or from *ex vivo* studies where cells harvested from the animals (typical spleen or lymph node cells), plated in cell culture and stimulated by inflammagens. Such primary and *ex vivo* cytokine analyses were considered separate records, as were cytokine analyses in different tissues. However, to avoid duplication, where the same cytokines were measured in the same source by two different methods (e.g. ELISA or qPCR) or at different time-points in the same cohort, only one was considered a record (specified in Supplementary Excel file).

### Data analysis

Based on the variables extracted, we determined that a formal meta-analysis was inappropriate due to substantial heterogeneity across studies^12^, including differences in disease models, helminth species, HDIPs, disease and cytokine measures, analytical methods and specific experimental design parameters. Instead, we conducted a cross-study quantitative analysis focusing on the percent change in disease burden and cytokine levels following treatment with helminth-derived peptides. To do so, data were extrapolated from the relevant figure(s) within each article using GetData Graph Digitizer software. The effects of the HDIPs were then quantitatively assessed by calculating the percent change in disease burden and cytokine levels in the animal models without and with HPID treatment. Data extraction was completed independently by two of the authors (SS and AF) and cross-checked for accuracy (ED).

### Statistical analyses

General study characteristics are shown as pie charts. All data related to percent disease or cytokine change are shown as scatter plots depicting individual records with the mean ± standard error of the mean (SEM). To determine if the HDIPs reduced or increased disease burden or cytokine levels beyond zero, data were analysed using one-sample t-test (with the hypothesised population mean set to zero). The reader is also directed to the Supplementary Excel File where the extracted data is provided in more detail and can be sorted, filtered and/or pivoted for further analyses.

## RESULTS

### General study characteristics

Sixty-five articles were included in this review in which HDIPs were assessed for disease-modifying and immunomodulatory effects in *in vivo* animal models of inflammatory disease (Table 1 and Supplementary Excel file). The HDIPs were assessed in multiple different disease models, but the most widely used were models of colitis (dextran sulphate sodium, trinitrobenzenesulfonic acid and T-cell transfer-induced), asthma (ovalbumin, *Alternaria* and house dust mite-induced), arthritis (collagen and monosodium urate-induced) and sepsis (lipopolysaccharide and caecal ligation/puncture-induced) (Fig. 2A). Other studies assessed HDIPs in animal models of diabetes, multiple sclerosis, pneumonia, rhinitis and bacterial infection and others as listed in Table 1. All models, with a single exception, were generated in mice. From these 65 articles, 141 records were extracted in which the disease-modifying effects of the HDIPs were assessed (94 clinical measures and 47 cellular measures), while 132 records assessed the effects of the HDIPs on cytokine levels (84 in primary sources and 48 in *ex vivo* analyses) (Fig. 2B).

**Fig. 2.**
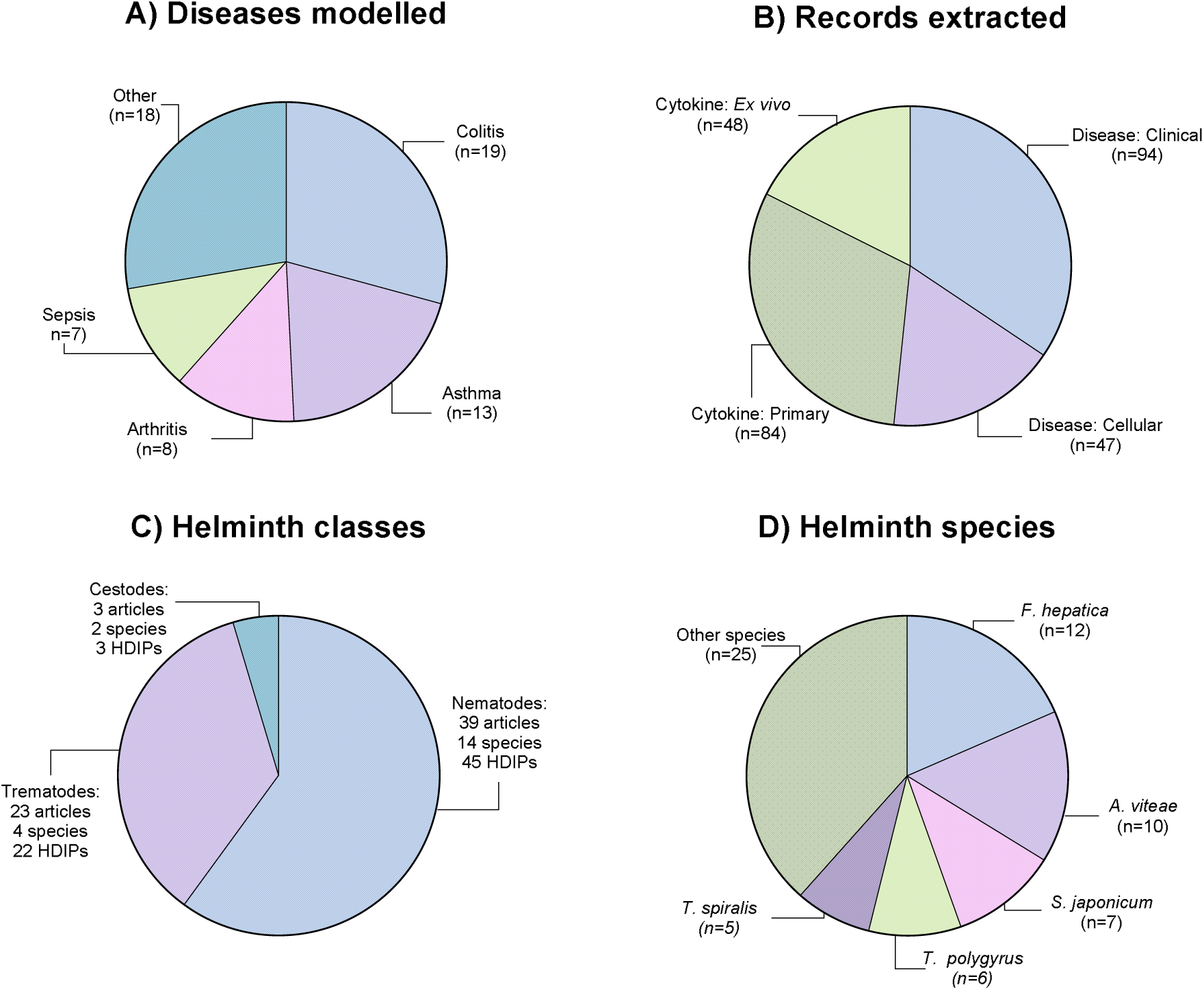
General study characteristics. A total of 65 articles were included in this review and a summary of the key characteristics of these articles is depicted in these pie charts. These show A) the main diseases modelled in these articles, B) the individual records extracted from these articles in which the HDIPs were assessed for effects on disease-modification or cytokine levels, C) the diversity of helminth classes, species and HDIPs assessed, and d) the most frequently assessed helminth species in these articles.

**Fig. 3.**
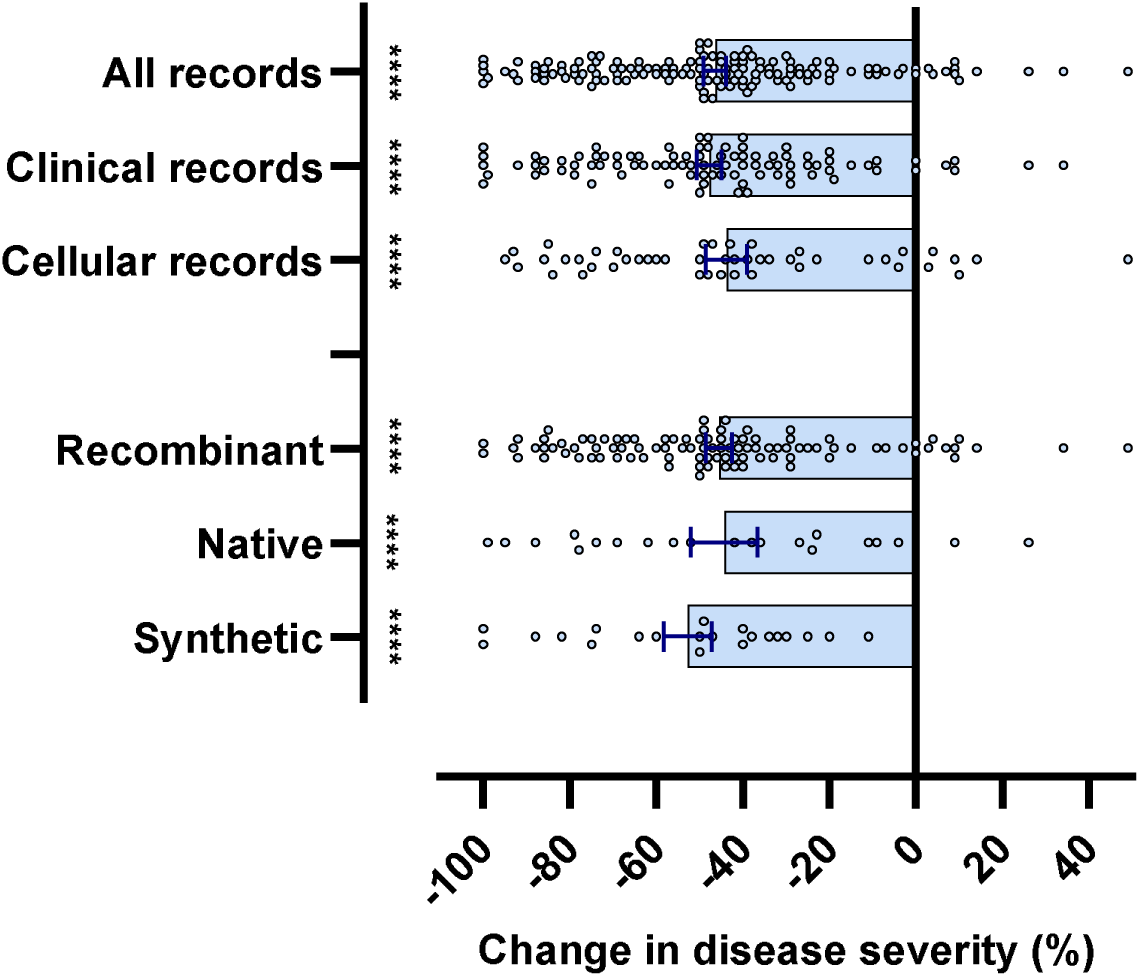
Effect of HDIPs on disease severity in the animal models. The HDIPs reduced disease severity across all disease models whether this was assessed using a clinical or a cellular measure, and whether the HDIP was native, recombinant or synthetic. Each data point represents a specific record extracted from Table 1/Supplementary Excel file, and the mean±sem is also shown. Data were analysed by one-sample test with the hypothesised population mean set to zero. \*\*\*\**P*<0.0001.

In the 65 selected articles, HDIPs from nematodes (roundworms) were most often tested, with 45 HDIPs from 14 species assessed in 39 articles (Fig. 2C). HDIPs from trematodes (flatworms/flukes) were also used frequently, with 22 HDIPs from 4 species assessed in 23 articles. In contrast, HDIPs from cestodes (flatworms/tapeworms) were only assessed in 3 articles. HDIPs from a total of 20 different helminth species were assessed in the selected studies. From these, the most widely used were HDIPs from the trematodes, *Fasciola hepatica* and *Schistosoma japonicum*, and HDIPs from the nematodes, *Acanthocheilonema viteae, Trichinella spiralis and Heligmosomoides polygyrus* (Fig. 2D). Most articles used recombinant forms of the HDIPs (64% of articles), while purified native HDIPs (22%) and synthetic HDIPs (13%) were less commonly used. From the different helminth species, >65 different HDIPs were tested, and although there was no widely used HDIP, ES-62 (a phosphorylcholine-containing glycoprotein secreted by *Acanthocheilonema viteae)* was used most frequently (n=9 articles).

### Effect of HDIPs on disease severity

From the 65 articles included in this review, 141 separate records were identified in which HDIPs were assessed for their effects on disease severity. This revealed that the HDIPs reduced the severity of disease across the range of inflammatory disease models assessed (All records: *t*_(140)_=18.30, *P*<0.0001). This was the case whether the disease severity was assessed using a clinical measure (Clinical records: *t*_(93)_=16.19, *P*<0.0001) or a cellular measure (Cellular records: *t*_(46)_=9.01, *P*<0.0001), and whether the HDIP was recombinant (Recombinant: *t*_(98)_=15.12, *P*<0.0001), native (Native: *t*_(20)_=5.73, *P*<0.0001) or synthetic (Synthetic: *t*_(20)_=9.43, *P*<0.0001). This was also the case with different routes of administration, different timings of administration (relative to disease induction) and different mechanisms of action of the HDIPs (Table 1).

This disease-modifying effect of the HDIPs was evident in the different inflammatory disease models assessed (Fig. 4A) including in colitis models (Colitis: *t*_(32)_=8.79, *P*<0.0001), asthma models (Asthma: *t*_(41)_=10.52; *P*<0.0001), arthritis models (Arthritis: *t*_(12)_=5.62, *P*<0.001), sepsis models (Sepsis: *t*_(9)_=6.29, *P*<0.001), as well as the other disease models assessed (Other: *t*_(42)_=8.91, *P*<0.0001), and was also common to HDIPs from all species used (Fig.4B) (*F. hepatica*: *t*_(19)_=6.11, *P*<0.0001; *A. viteae*: *t*_(20)_=5.11, *P*<0.0001; *S. japonicum*: *t*_(16)_=6.15, *P*<0.0001; *H. polygyrus*: *t*_(11)_=5.02, *P*<0.001; T. spiralis: *t*_(10)_=6.17, *P*<0.001; other species: *t*_(54)_=12.72, *P*<0.0001).

**Fig. 4.**
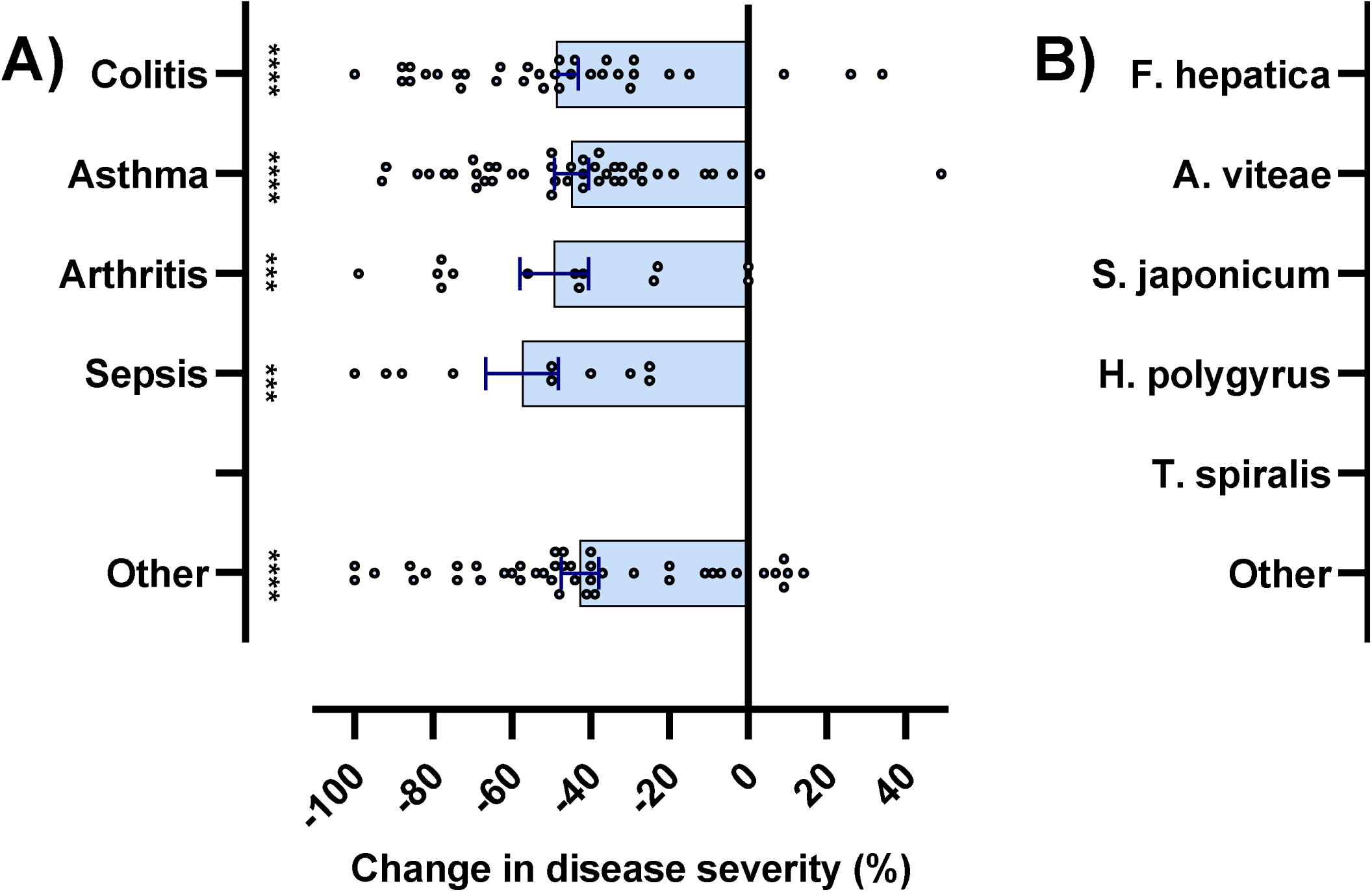
Effect of HDIPs on disease severity in the animal models. The HDIPs reduced disease severity across all disease models assessed, and this was common to all helminth species. Each data point represents a specific record extracted from Table 1/Supplementary Excel file, and the mean±sem is also shown. Data were analysed by one-sample test with the hypothesised population mean set to zero. \*\*\*\**P*<0.0001, \*\*\**P*<0.001.

### Effect of HDIPs on cytokines

From the 65 articles included in this review, 132 records assessed the effects of the HDIPs on cytokine levels in animal models of inflammatory disease. In the initial analysis, the cytokines, TNFα, INFγ, Il-1β, IL-5, IL6, IL12 and IL-17, were grouped together as pro-inflammatory cytokines, while TGFβ, IL-4, IL-10 and IL-13 were grouped as anti-inflammatory cytokines. The HDIPs reduced the levels of these pro-inflammatory cytokines across the animal models of inflammatory diseases (Fig. 5A) (All records: *t*_(253)_=13.38, *P*<0.0001), and this was the case whether the cytokines were measured in primary sources from the animals (Primary sources: *t*_170)_=14.76, *P*<0.0001) or in *ex vivo* studies using cells harvested from HDIP-treated animals (*Ex vivo* sources: *t*_(82)_=5.57, *P*<0.0001), and whether the HDIP was recombinant (Recombinant: *t*_(172)_=15.22, *P*<0.0001) or native (Native: *t*_(50)_=6.72, *P*<0.0001). Overall, the synthetic HDIPs did not significantly reduce the levels of pro-inflammatory cytokines (Synthetic: *t*_(29)_=0.85, *P*<0.0001). However, the HDIPs did increase levels of these anti-inflammatory cytokines across the animal models of inflammatory diseases (Fig. 5B) (All records: *t*_(188)_=4.68, *P*<0.0001), and this was the case whether the cytokines were measured in primary or *ex vivo* sources (Primary sources: *t*_136)_=5.06, *P*<0.0001; *Ex vivo* sources: *t*_(51)_=2.49, *P*<0.05), and whether the HDIP was recombinant (Recombinant: *t*_(137)_=4.93, *P*<0.0001) or synthetic (Synthetic: *t*_(29)_=2.99, *P*<0.01). Overall, the native HDIPs did not significantly increase the levels of pro-inflammatory cytokines (Native: *t*_(20)_=1.34, *P*>0.05).

**Fig. 5.**
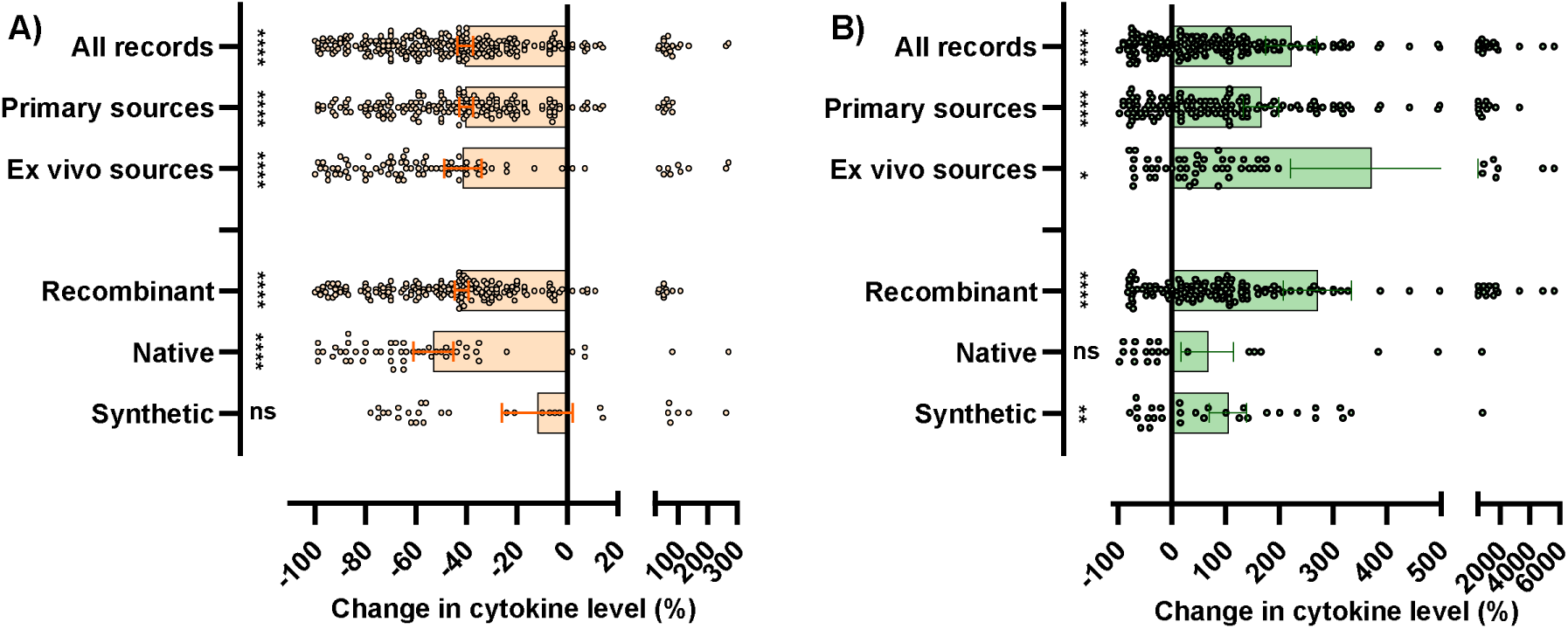
Effect of HDIPs on cytokine levels in the animal models. A) With the exception of the synthetic forms, the HDIPs reduced pro-inflammatory cytokine levels in the disease models whether this was measured in primary sources or *ex vivo*, and whether the HDIP was recombinant or native. B) With the exception of the native forms, the HDIPs increased anti-inflammatory cytokine levels in the disease models whether this was measured in primary sources or *ex vivo*, and whether the HDIP was recombinant or synthetic. Each data point represents a specific record extracted from Table 1/Supplementary Excel file, and the mean±sem is also shown. Data were analysed by one-sample test with the hypothesised population mean set to zero. \*\*\*\**P*<0.0001, \*\**P*<0.01, \**P*<0.05, ns: not-significantly different to zero.

This ability of the HDIPs to reduce pro-inflammatory cytokine levels was a property of HDIPs from most species used (Fig.6A) (*F. hepatica*: *t*_(59)_=4.39, *P*<0.0001; *A. viteae*: *t*_(26)_=2.46, *P*<0.05; *S. japonicum*: *t*_(78)_=6.60, *P*<0.0001; *H. polygyrus*: *t*_(5)_=1.92, *P*>0.05; *T. spiralis*: *t*_(49)_=4.97, *P*<0.0001; other species: *t*_(82)_=15.10, *P*<0.0001). Similarly, the HDIPs from most species were also capable of increasing anti-inflammatory cytokine levels (Fig.6B) (*F. hepatica*: *t*_(12)_=1.58, *P*>0.05; *A. viteae*: *t*_(19)_=1.21, *P*>0.05; *S. japonicum*: *t*_(41)_=2.30, *P*<0.05; *H. polygyrus*: too few points; *T. spiralis*: *t*_(53)_=4.17, *P*<0.001; other species: *t*_(83)_=4.01, *P*<0.001).

**Fig. 6.**
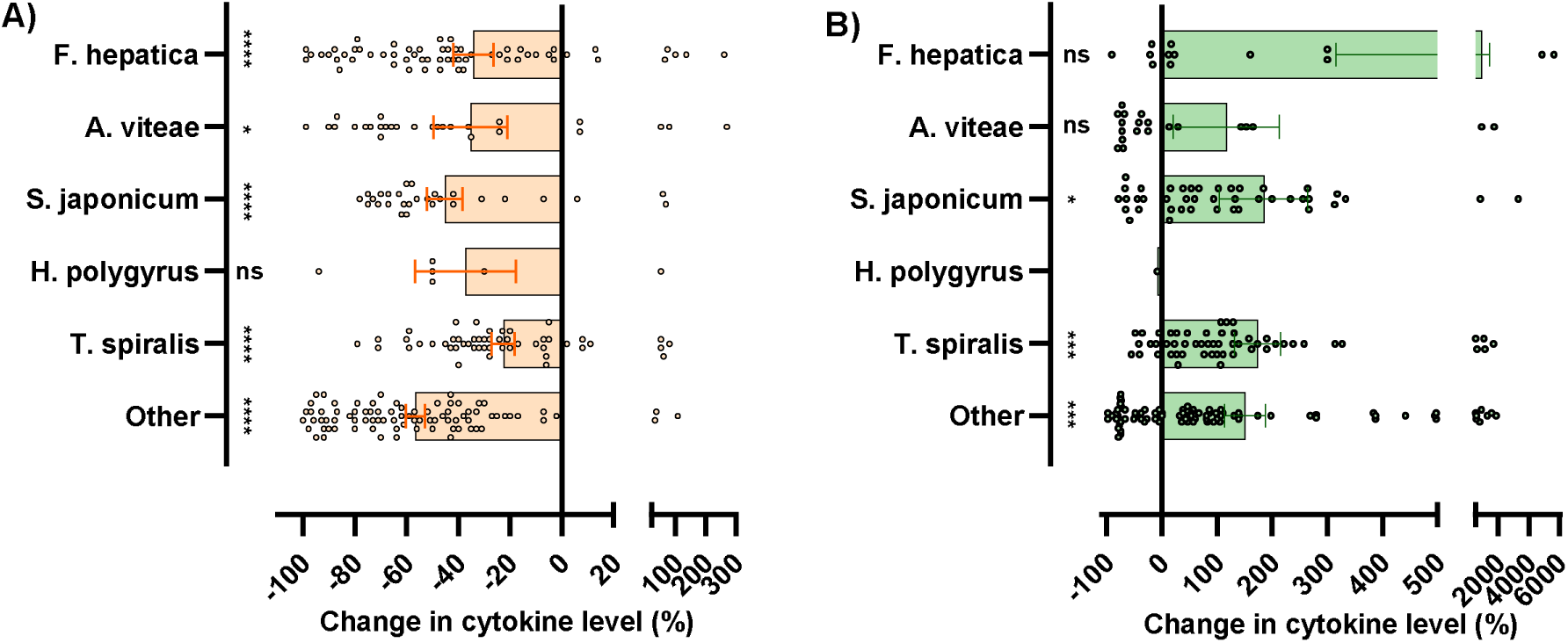
Effect of HDIPs on cytokine levels in the animal models. A) With the exception of *H. polygyrus*, the HDIPs from all species of helminth reduced pro-inflammatory cytokine levels in the disease models. B) With the exception of *F. hepatica* and *A. vitae*, the HDIPs from all species of helminth increased anti-inflammatory cytokine levels in the disease models. Each data point represents a specific record extracted from Table 1/Supplementary Excel file, and the mean±sem is also shown. Data were analysed by one-sample test with the hypothesised population mean set to zero. \*\*\*\**P*<0.0001, \*\*\**P*<0.001, \**P*<0.05, ns: not-significantly different to zero.

When the effects of HDIPs on individual cytokines was assessed, this revealed that they reduced levels of all pro-inflammatory cytokines measured, except for IL-5 (Fig. 7A) (Il-1β: *t*_(15)_=9.27, *P*<0.0001; TNFα: *t*_(60)_=11.16, *P*<0.0001; IL6: *t*_(42)_=6.77, *P*<0.0001; INFγ: *t*_(64)_=6.38, *P*<0.0001; IL12: *t*_(13)_=3.86, *P*<0.01; IL-17: *t*_(30)_=3.04, *P*<0.01; IL-5: *t*_(23)_=1.66, *P*>0.05). With regard to the anti-inflammatory cytokines, the HDIPs increased levels of IL-10 and TGFβ but not IL-4 or IL-13 (Fig. 7B) (Il-10: *t*_(81)_=3.73, *P*<0.001; TGFβ: *t*_(40)_=4.39, *P*<0.0001; IL-4: *t*_(43)_=1.20, *P*>0.05; Il-13: *t*_(21)_=0.03, *P*>0.05).

**Fig. 7.**
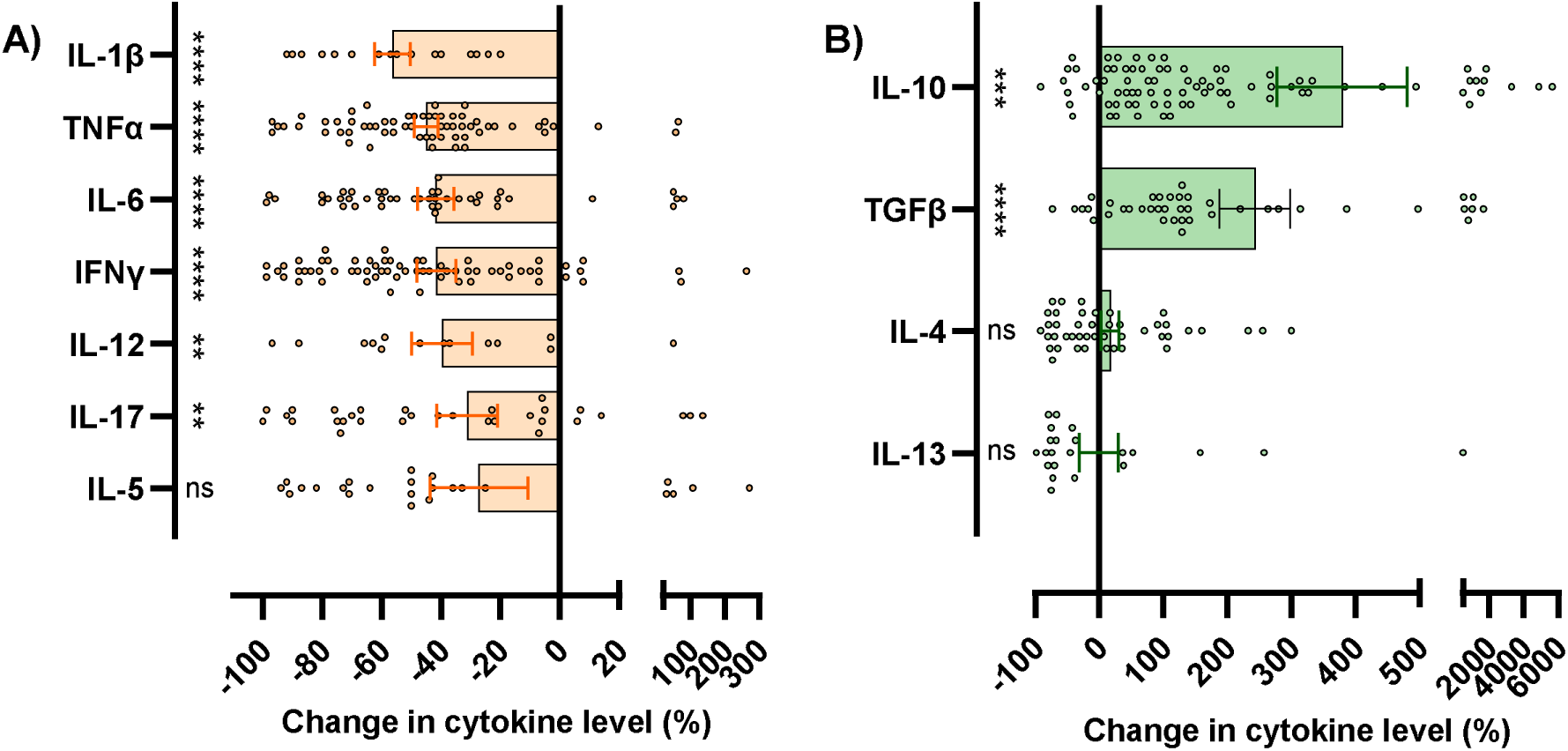
Effect of HDIPs on individual cytokine levels in the animal models. A) With the exception of IL-5, the HDIPs reduced all pro-inflammatory cytokines in the disease models. B) The HDIPs increased levels of IL-10 and TGFβ, but not IL-4 or IL-13, in the disease models. Each data point represents a specific record extracted from Table 1/Supplementary Excel file, and the mean±sem is also shown. Data were analysed by one-sample test with the hypothesised population mean set to zero. \*\*\*\**P*<0.0001, \*\*\**P*<0.001, \*\**P*<0.01, ns: not-significantly different to zero.

## Discussion

Helminths have long been recognized for their potent immunomodulatory capacity, enabling chronic persistence within immunocompetent hosts through active manipulation of selective host immune pathways^4–7^. While early translational efforts focused on live helminth therapy, concerns regarding safety, variability in host responses, and regulatory feasibility have shifted attention toward using defined helminth-derived immunomodulatory products. In particular, isolated HDIPs offer a more controlled and potentially safer alternative, with reduced risk of infection and off-target effects compared to live organisms. A growing number of HDIPs, originating from diverse helminth species, have been characterized with distinct mechanisms of action and evaluated in both *in vitro*^10^ and *in vivo* animal models of inflammatory and autoimmune disease. Despite this expanding preclinical body of evidence, clinical translation remains limited, with only a single HDIP advancing to clinical trial evaluation^9^. This translational bottleneck underscores the need for systematic assessment of the existing *in vivo* data to better define efficacy and therapeutic potential. To address this gap, the present study systematically reviews the *in vivo* animal models in which HDIPs have been tested, with particular emphasis on their capacity to attenuate disease activity and modulate inflammatory cytokine responses.

Colitis, asthma, arthritis and sepsis were the most frequently investigated disease models, although additional inflammatory conditions - including diabetes, multiple sclerosis, pneumonia, rhinitis and bacterial infection - were also examined. Experimental colitis was induced using established chemical models such as trinitrobenzene sulfonic acid and dextran sulphate sodium, as well as adoptive T-cell transfer approaches. These models produce a reproducible clinical phenotype characterized by weight loss, altered stool consistency, and rectal bleeding, typically quantified using the Disease Activity Index (DAI^78^). Regardless of the colitis model employed, the helminth species of origin, the specific HDIP administered, or whether the peptide was recombinant, native or synthetic, HDIP treatment uniformly resulted in a significant reduction in disease severity.

A similar pattern was observed in the other inflammatory models. Asthma was induced using ovalbumin, *Alternaria*, or house dust mite allergens; arthritis was modelled using collagen-induced or monosodium urate-induced inflammation; and sepsis was triggered via lipopolysaccharide administration or caecal ligation and puncture. Across all of these models - as well as in the less frequently studied disease settings - HDIP administration consistently led to significant improvements in disease outcomes. These improvements were demonstrated through reductions in clinical, macroscopic or histopathological scores, or through functional measures specific to each model. Collectively, these findings highlight the broad and reproducible anti-inflammatory efficacy of HDIPs across diverse animal models of human inflammatory disease.

In parallel with the reduction in disease severity, HDIPs treatment was also associated with a marked reduction in pro-inflammatory cytokines, alongside increased production of anti-inflammatory cytokines. In the selected articles, cytokines were measured in two broad sources: 1) primary sources in which the cytokines were measured directly in fluids (e.g. serum, bronchiolar lavage), cells (e.g. splenocytes) or tissues (e.g. colon, lung) from the animals models; and 2) secondary sources where cells (usually from the spleen or lymph nodes) were harvested from the HDIP-treated animal models, plated in cell culture and assessed *ex vivo* for cytokine responses (usually after stimulation with an inflammagen). In both of these sources, regardless of the helminth species or specific HDIP, the peptides significantly reduced levels of pro-inflammatory cytokines including IL-1β, TNFα, IFNγ and IL-6, and increased levels of the anti-inflammatory cytokines, TGFβ and IL-10. Interestingly, some differences between recombinant, native and synthetic HDIPs emerged here. Specifically, the synthetic forms did not significantly reduce pro-inflammatory cytokine levels, while the native forms did not significantly increase anti-inflammatory cytokine levels. However, as these were assessed far less frequently than recombinant forms, it is likely that this simply reflects the smaller sample size.

A notable feature of the literature is the near-exclusive reliance on murine models. With only one study conducted in rats, the evidence base for therapeutic efficacy remains heavily dependent on a single species. While mouse models undoubtedly provide valuable insight, broader validation in additional animal models may strengthen the preclinical evidence base and enhance confidence in human translation. Another striking aspect of the literature selected through this systematic screen is the paucity of studies focusing on neuroinflammatory and/or neuroimmune conditions. Even through neuroinflammation is a central feature, and even a driving force, in many neurodegenerative diseases^79^, the potential therapeutic efficacy of HDIPs for such conditions has not yet been explored. Given that HDIPs show efficacy across a range of diverse and mechanistically-distinct inflammatory disease models, this highlights the need to investigate their potential across a range of inflammatory related diseases, including neurological ones. Four articles did assess HIPDs in murine models of multiple sclerosis^36,43,73,75^, and these (mostly) demonstrated disease-modifying potential, thus providing some rationale for assessment in other neurological conditions in which there is an inflammatory/immune component.

Taken together, the evidence supports a conclusion that HDIPs exert broad, reproducible disease-modifying effects *in vivo* across multiple inflammatory pathologies. These effects are characterised by significant reductions in clinical and histopathological disease severity, underpinned by suppression and enhancement of key pro-inflammatory and anti-inflammatory cytokines respectively. HDIPs from 20 helminth species were evaluated, spanning nematodes, trematodes, and cestodes. Despite this taxonomic diversity, the capacity to reduce disease severity was a shared feature across species supporting the concept that helminth-derived molecules represent a rich and evolutionarily refined source of immunoregulatory therapeutics. Viewed collectively, the studies underpinning this systematic review encourage accelerated advancement of HDIPs to first-in-human clinical trials.

## Data availability

All data is available in the Supplementary Excel file.

## Author Contributions

SS, AF and ED led the design and implementation of the systematic review, and wrote the first draft of the manuscript; RL, SD, JPD and DMK provided expert input into the drafts of the manuscript.

